# New forms of ecological knowledge acquisition for freshwater mussels’ conservation

**DOI:** 10.1101/2022.09.06.506734

**Authors:** Noé Ferreira-Rodríguez

## Abstract

Freshwater mussels are of recognized importance for ecosystem functioning, for the provision of ecosystem services, and for human well-being. Whether people are more concerned about freshwater mussels has a large role to play in promoting and supporting political decisions that protect their habitat. A pilot study was performed in the Midwest of the United States to evaluate the awareness people have about freshwater mussels both native and non-native. Using semi-structured face-to-face interviews, I explore whether ecological knowledge is still in place in the community, and what the relationship is between knowledge, age and urbanization. Multinomial logistic regression has shown that population size and age class are significant predictors of ecological knowledge. Present results suggest that common channels of ecological knowledge transmission have been interrupted long ago. As a result, local ecological knowledge is being substituted by modern ecological knowledge. Nevertheless, new processes of ecological knowledge acquisition emerge. Under this scenario, future management prospects would rely on state institutions which are implementing alternative dissemination pathways in modern socio-ecological systems.

## Introduction

The Millennium Ecosystem Assessment defines existence value as the “value to knowing that a resource exists” (Millennium Ecosystem Assessment 2005). This intangible valuation is related with the amount an individual would pay to know that a particular species exists in its natural habitat (Loomis et al. 2000). However, before anyone can be willing to pay for a given ecosystem service, they should be aware of their existence.

Unionid mussels, a dominant part of freshwater biodiversity, are key components of ecosystems, providing a wide array of services useful for human well-being (Vaughn 2018). Freshwater mussels are, however, one of the most threatened animal groups (Geist 2011). Evidences of their decline came from the 1970s, when the first extinctions were documented in North America (Stansbery 1971). A variety of factors, including habitat alteration, invasive species and climate change seems to be the primary drivers of their poor conservation status (Ferreira-Rodríguez et al. 2019). Hence, as freshwater mussels decline, also their role in ecosystem functioning is at risk.

Accumulated knowledge about ecosystems and their biodiversity is an important part of people’s capacity to manage and conserve the environment, and has a large role to play in decision-making (Cullen et al. 2007). But, this ecological knowledge is being rapidly lost in modern – wealthier, urbanized, and globalized – societies (Pilgrim et al. 2008). Under this scenario, the first question is whether ecological knowledge is still in place in the community, and what the relationship is between knowledge, age and urbanization. In addition, the zebra mussel *Dreissena polymorpha* (Pallas, 1771) has been introduced and expanded its range from the Great Lakes to Oklahoma between 1986 and 1993 (http://nas.er.usgs.gov; accessed September 30, 2020). Given the high economic and ecological costs associated to *D. polymorpha* invasions, efforts are underway to increase awareness among population about the harmful effects of this invasive species to ecosystems and human wellbeing. Hence, the second question is whether ecological knowledge is reoriented towards invasive species at the expense of natives. In order to explore these questions, a pilot study using semistructured face-to-face interviews has been carried out in the Midwest of the United States.

## Material and methods

This work was performed in Minnesota and Kansas between May and June 2018. A stratified cluster sampling design was used for the selection of the survey sample. First, cluster sampling was used to group population nucleuses within each state in terms of three population levels (>10 000; 10 000-100 000; and <100 000 inhabitants). Second, stratified and aleatory (at random) sampling was used to ensure that population distribution within the three population levels was fairly represented. The sampling strategy included stakeholders residing in or visiting the area. Interviewees were randomly selected from populated areas including gas stations, community-buildings, supermarkets, and recreational areas, among others. We had no contact with any of the interviewees in advance of our surveys. The sampling frame was restricted to individuals over 18 years old, and interviewees were grouped in three age classes (age estimation based on physical appearance); i.e. young adulthood 18-40, middle-age 41-60, and old adulthood 61 and over. A total of 71 face-to-face surveys at seven population nucleuses (4 in Minnesota, 3 in Kansas) were conducted. In the light of interviewees’ attitude (i.e. bad, not so good, good), and subsequent doubts about the validity of responses, we restricted the starting dataset to interviewees that provided reliable information; i.e. respondents with “bad” and “not so good” attitude [i.e., uncooperative behaviour, negatively worded items; *sensu* Stocké and Langfeldt (2004); n = 4] were excluded from the data analysis.

Interviewees were asked to respond to a survey on a voluntary basis and those who participated were not coerced into responding to questions. We informed them that all responses were anonymous, that we just wanted to know their opinion, and that there were no right answers. The questions were related to knowledge about freshwater mussels (Supplementary material 1). The answers were linked to socio-demographic information of the respondents. We used descriptive analysis to characterize knowledge about freshwater mussels’ existence. Multinomial logistic regressions were used to elucidate the relationships between two independent variables (population size and age class) and one nominal dependent variable (freshwater mussels’ knowledge). In a second analysis, two nominal dependent variables (native unionid and non-native dreissenid mussels’ knowledge) were used. All statistical tests were conducted using IBM SPSS Statistics Version 19 (SPSS, Chicago, IL).

## Results

Half of the interviewees (50.8%) gave a positive answer to the question about freshwater mussels’ existence. Among them, 28.4% of the interviewees referred to unionid mussels, and 22.4% to dreissenid mussels.

The proposed explanatory model (Table 1) as a whole fitted significantly better than the empty model (i.e. model with no predictors) to the data (χ^2^ = 17.622, d.f. = 8, p = 0.024). Specifically, tests for the main effect of population size (χ^2^ = 11.248, d.f. = 4, p = 0.024) was statistically significant at significance level 5 percent, and age class (χ^2^ = 8.420, d.f. = 4, p = 0.077) was statistically significant at significance level 10 percent.

**Table 1.**
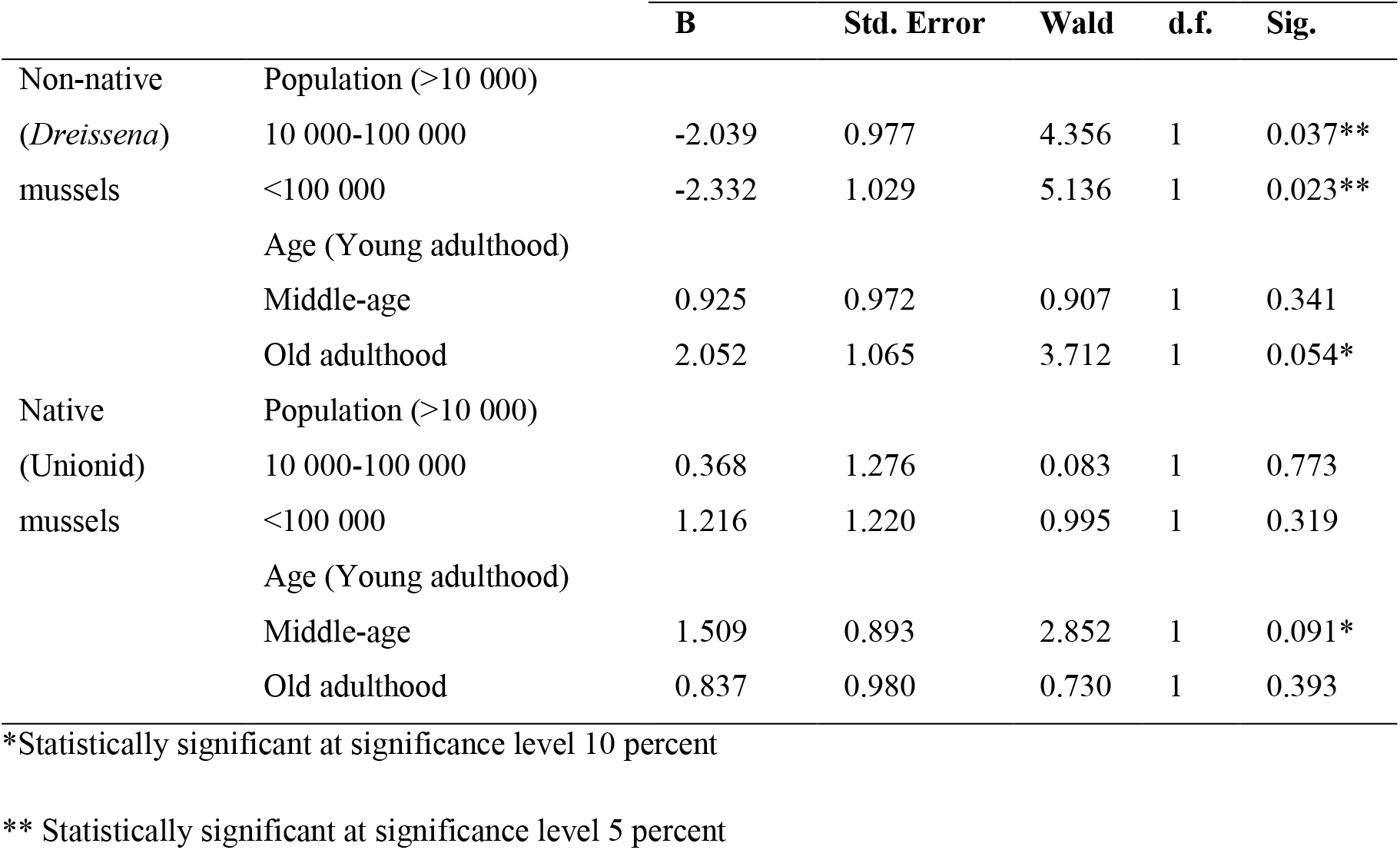
Predictors of freshwater mussels’ knowledge from adjusted multinomial logistic regression model (n = 67). Knowledge about native freshwater mussels (Unionidae) and non-native freshwater mussels (*Dreissena polymorpha*) existence, with respect to the reference category not knowledge about freshwater mussels’ existence.

|  |  | **B** | **Std. Error** | **Wald** | **d.f.** | **Sig.** |
| --- | --- | --- | --- | --- | --- | --- |
| Non-native<br>(*Dreissena*)<br>mussels | Population (>10 000) |  |  |  |  |  |
|  | 10 000-100 000 | -2.039 | 0.977 | 4.356 | 1 | 0.037** |
|  | <100 000 | -2.332 | 1.029 | 5.136 | 1 | 0.023** |
|  | Age (Young adulthood) |  |  |  |  |  |
|  | Middle-age | 0.925 | 0.972 | 0.907 | 1 | 0.341 |
|  | Old adulthood | 2.052 | 1.065 | 3.712 | 1 | 0.054* |
| Native<br>(Unionid)<br>mussels | Population (>10 000) |  |  |  |  |  |
|  | 10 000-100 000 | 0.368 | 1.276 | 0.083 | 1 | 0.773 |
|  | <100 000 | 1.216 | 1.220 | 0.995 | 1 | 0.319 |
|  | Age (Young adulthood) |  |  |  |  |  |
|  | Middle-age | 1.509 | 0.893 | 2.852 | 1 | 0.091* |
|  | Old adulthood | 0.837 | 0.980 | 0.730 | 1 | 0.393 |

\*Statistically significant at significance level 10 percent
\*\* Statistically significant at significance level 5 percent

The relative log odds of finding one person aware of unionid mussels’ existence versus finding one person with no knowledge about freshwater mussels existence increased by 1.509 if asking to one middle-aged person versus asking to young adulthood person. The relative log odds of finding one person aware of dreissenid mussels’ existence versus finding one person with no knowledge about freshwater mussels’ existence increased by 2.052 if asking to old adulthood person versus asking to young adulthood person. The relative log odds of finding one person aware of dreissenid mussels’ existence versus finding one person with no knowledge about freshwater mussels existence at the significance level 5 percent decreased by 2.039 and by 2.332 if moving from small population nucleuses (< 10 000 habitants) to large ones (>100 000 habitants).

## Discussion

Although still preliminary, present results have shown that in the older age class and those residing in rural areas are more aware about dreissenid mussels’ existence than young adulthood ones and those residing in large population nucleuses. In this regard, ecological knowledge acquisition is based on an accumulation of observations and saturation occurs at an elderly age in wealthier countries (Pilgrim et al. 2008). Hence, it is not surprisingly that old adulthood groups will be more aware about mussels’ existence than younger ones. In addition, resource use practice is reflected in ecological knowledge. As a consequence, in rural areas, the proximity of users to the resource confers the ability to observe day-to-day changes in the system (Berkes and Turner 2006). Concerning unionid mussels’ existence, results have shown the absence of differences between old and young adulthood groups. These results suggest that ecological knowledge could have been lost long time ago (>70 years). In this vein, freshwater mussels harvesting for the pearl industry was common until 1950s. Afterwards and despite sporadically reported by some interviewees (results not shown), these practices have fallen out because they have been now very limited or completely banned (Thorp and Rogers 2010; Vaughn 2018). However, more qualitative data are needed to precisely determine when knowledge was acquired and lost.

The most common channels of ecological knowledge transmission among generations (i.e. through observations and narratives) seems to be interrupted with the shift from rural to urban societies. Under this scenario, new forms of knowledge acquisition and learning emerge. As an example, environmental education programs play a vital role in increasing social awareness towards freshwater mussels. Among them, the implementation of educational and captive breeding programs by the American Zoos and Aquariums Association (AZA Association) provides a national wide example of new forms of knowledge acquisition in urban areas. The AZA Association has prompted increasing efforts to disseminate the role of freshwater mussels in the provisioning of ecosystem services among the population through a collaborative network of institutions across the country. However, ecological knowledge is a cumulative body of knowledge evolving by adaptive processes. In this sense, knowledge acquisition through the accommodation of new forms of information may be the modern versions of traditional ecological knowledge transference among generations. As main receivers of these new dissemination pathways, middle-age people were more aware than young adulthood people about freshwater mussels’ existence. However, the real impact of this kind of environmental educational programs on public awareness remains a matter of future research.

## Conclusions

The recognition of the important ecosystem services – other than provisioning ecosystem services – provided by unionid mussels, led to a more conservationist approach in wealthier socio-ecological systems. It should be stressed, however, that traditional ecological knowledge about unionid mussels is likely to be substituted by modern ecological knowledge about invasive species, especially among old adulthood groups and in rural areas. Therefore, future management prospects would rely on state institutions which are implementing alternative dissemination pathways for increasing people’s awareness about unionid mussels as part of the conservation strategy not only in urbanized areas but in rural areas as well. Despite the results suggest the utility of the ongoing educational programs, their effectiveness should be empirically validated and widely implemented for increasing public awareness and, in consequence, the existence value of freshwater mussels with conservation purposes.

## Acknowledgements

I am very grateful to Daniela Irimia for her unconditional support performing the personal interviews and who is largely responsible for its successful outcome. I also am very grateful to Ben Minerich from the Minnesota Zoo (Apple Valley, Minnesota) for his kindly support performing the field surveys. I also want to thank Andy Allison from the National Mississippi River Museum & Aquarium (Dubuque, Iowa) and Mike Brittsan from the Columbus Zoo and Aquarium (Powell, Ohio) for their willingness to help with this research. I am very grateful to the Oklahoma Biological Survey (University of Oklahoma, Norman, Oklahoma) and especially to Professor Caryn C. Vaughn for providing a quiet place for working in the USA. I would like to thank Antonio Castro for some discussions that helped shape the research. The questionnaire was designed following the example of previous studies approved by the University of Oklahoma Institutional Review Board for the Protection of Human Participants (permit number 2733). I was supported by a post-doctoral fellowship (Xunta de Galicia Plan I2C 2017-2020, 09.40.561B.444.0) from the government of the autonomous community of Galicia.

