## Supplementary material 1 for "New forms of ecological knowledge acquisition for freshwater mussels’ conservation"

DATE ............................ Nº SURVEY ………………………………. PLACE ……………………………….

GENDER ………………………………. ETHNIC BACKGROUND ……………………………….

AGE ☐ 18-40 years ☐ 40 – 60 years ☐ > 60 years

Respondent's attitude: good/not very interested/not interested

Respondent's understanding: good/not very interested/not interested

We are studying the general knowledge about freshwater mussels. To do this we are surveying locals and tourists in the area. It would be helpful to know your opinion/perception through this survey. Remember that all responses are anonymous and it only takes a few minutes. There are no "right answers", just tell us what you think. Thank you for your help!

**Study area**

1. Where do you live? ………………………………………… (Town)

2. How often do you visit rivers or lakes? …………………………………………

**Freshwater mussels' knowledge**

1. - Do you know what freshwater mussels are? ☐ NO ☐ YES

2. - Do you know any problems related to freshwater mussels? ☐ NO ☐ YES

If yes specify?……………………………….

*For the interviewer: Is she/he talking about Unionid mussels/Dressenid mussels

3. - Do you known any benefit related freshwater mussels?? ☐ NO ☐ YES

If yes specify?………………………………. (e.g. food for humans, water purification, pearls)
